# Target isoforms are an overlooked challenge and opportunity in chimeric antigen receptor cell therapy

**DOI:** 10.1101/2022.03.02.482604

**Authors:** Mike Bogetofte Barnkob, Kristoffer Vitting-Seerup, Lars Rønn Olsen

## Abstract

The development of novel chimeric antigen receptor (CAR) cell therapies is rapidly growing, with 299 new agents being reported and 109 new clinical trials initiated so far this year. One critical lesson from approved CD19-specific CAR therapies is that target isoform switching has been shown to cause tumor relapse, but little is known about the isoforms of CAR targets in solid cancers. Here we assess the protein isoform landscape and identify both the challenges and opportunities protein isoform switching present as CAR therapy is applied to solid cancers.

## Main text

Isoforms arise when different exons are combined through RNA splicing, and are translated into proteins with distinct properties. Especially in cancer cells, dysregulation and alternative splicing is thought to be involved in many hallmarks of cancer^1^. At least 75% of human protein-coding genes give rise to multiple distinct protein isoforms^2^, which can severely affect the sensitivity of therapeutic targeting of specific proteins. In a recent study of 883 small molecule cancer drugs targeting 1,434 different proteins, the authors found that 76% of these drugs would miss a target isoform if a switch occurred, or induce off-target effects in isoforms expressed in normal tissues^3^. Because isoforms of a protein may be differentially expressed in individual cancer cells^4^, they may produce a pool from which escape variants can arise when selective pressure is exerted by targeted therapy. Such adaptive resistance to targets has been seen when using CAR T cells to target EGFRvIII in glioblastoma multiforme^5^ and CD19 in B-cell acute lymphoblastic leukemia^6–8^

In order to assess the level of isoform switching in tumors, we analyzed RNA-seq data from TCGA and compared 5,562 tumor samples spanning 12 solid cancers to matched healthy tissue (see **Supplementary Methods**). As previously shown^9^, protein isoform switches are very frequent in cancers (**Figure 1A**). Such isoform switches greatly impact the sequences of the expressed proteins (**Figure 1B**), which can negatively affect the targetable epitopes of tumor-associated antigens. In fact, the challenge is even more pertinent for antibody-based therapies, such as CAR cell therapy: our analyses showed that isoform switches in cell membrane proteins are significantly enriched in 8 of the 12 cancers investigated (**Figure 1C**), with an average of 141 cell membrane proteins affected per cancer type.

**Figure 1.**
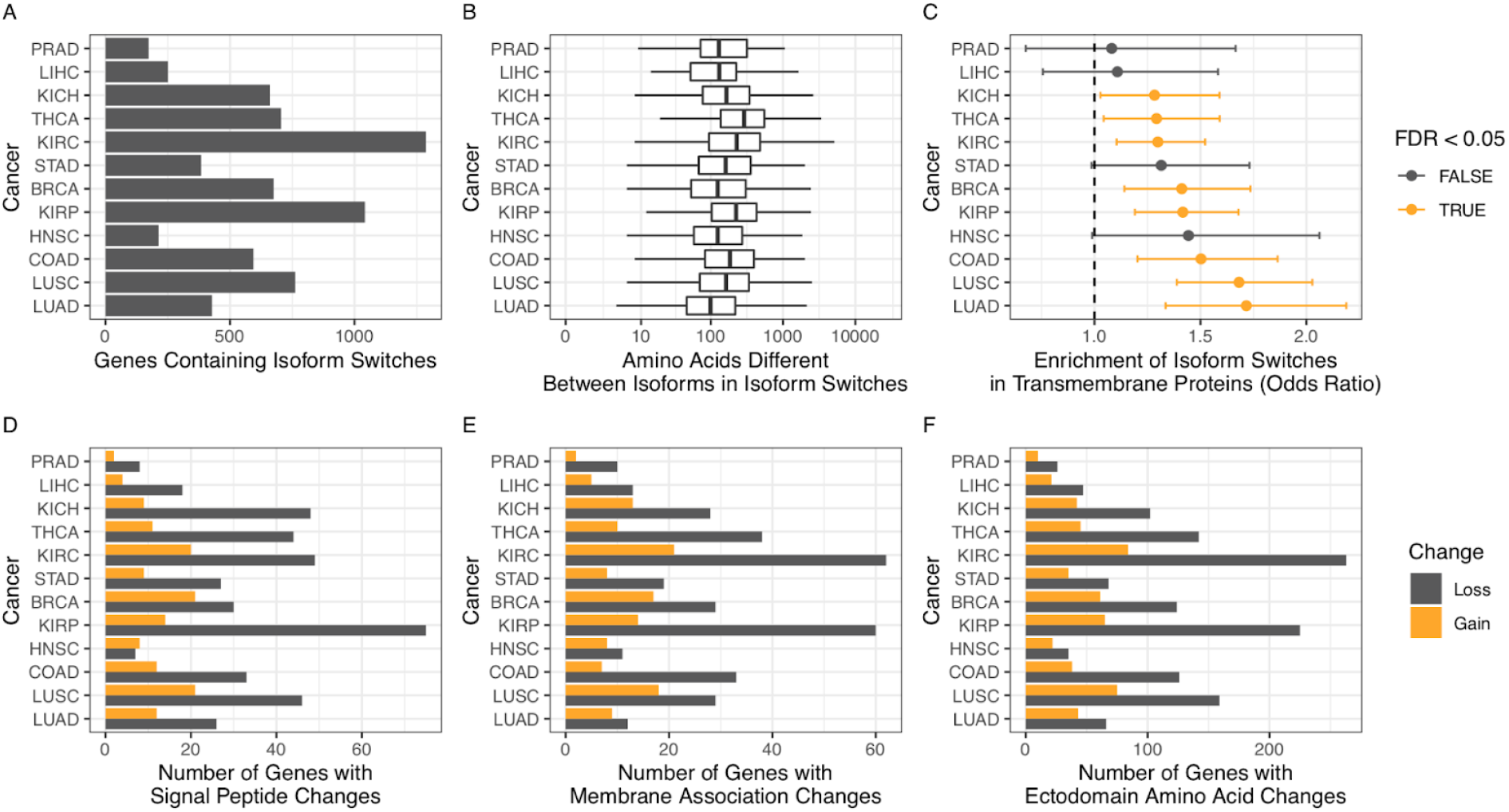
Protein isoform switching frequently occurs in tumor tissue. (a) The number of genes containing at least one isoform switch in 12 solid cancer types (b) Boxplot showing the number of amino acids being different between protein isoforms in each isoform switch. Outliers are not shown. (c) The enrichment of isoform switches in cell membrane proteins. Enrichment is given as odds-ratios (dot) along with 95% confidence interval (error bar). Color denotes false discovery rate (FDR) corrected P-values < 0.05. (d) Number of genes where a signal peptide is gained or lost (as denoted by color) due to isoform switches. (e) Number of genes where an isoform gain or loss (as denoted by color) membrane association. (f) Number of genes where amino acids in the ectodomain are gained or lost (as denoted by color) due to isoform switches.

The enrichment of isoform switches in cell membrane proteins compared to intracellular proteins prompted us to more thoroughly analyze both the opportunities and risks that isoform switches present. We first examined changes in signaling peptides of membrane-associated proteins as these play an important role in determining the subcellular localization and secretion status of the protein^10^. Across all 12 solid cancers, 242 membrane proteins lost signaling peptides (**Figure 1D**), indicating that these isoforms would no longer be secreted and thus be more amenable for targeting. However, our analysis also showed that 180 proteins not only lost their membrane association (**Figure 1E**), but 710 of those that remained membrane-bound, lost portions of their ectodomains (**Figure 1F**). Jointly this shows that many isoforms used in solid cancers are less targetable than their normal tissue counterparts. Interestingly, we also identified a small, but consistent number of genes in cancer cells that either gained membrane association or additional ectodomain amino acids (**Figure 1E-F**) indicating that isoform switches also lead to changes that could be potential tumor-specific targets for CAR therapy.

To better understand how isoforms might affect the success of new CAR therapies, we next examined the isoform status of the top five membrane proteins currently being tested in clinical CAR trials against solid tumors. These are the carcinoembryonic antigen-related cell adhesion molecule 5 (CEA; *CEACAM5),* Mucin-1 ( *MUC1),* Glypican-3 ( *GPC3),* Mesothelin ( *MSLN),* and receptor tyrosine-protein kinase erbB-2 (HER2; *ERBB2)*^11^.

We compared the isoforms expressed in the targeted cancer types and corresponding healthy tissue. The number of isoforms ranged from 4 to 23 for the five genes (**Figure 2A, Supplementary Figure 1A, 2A, 3A and 4A**), all with different expression levels between normal tissue and cancer (**Figure 2B-D; Supplementary Figure 1B, 2B-D, 3B and 4B**). GPC3 was unique in having isoforms that were all associated with the cell membrane, while the other four targets had isoforms that were either secreted, associated with intracellular membranes, the cytoplasm, or the cell membrane (**Figure 2E; Supplementary Figure 1C, 2E, 3C and 4C**). As indicated by our analysis across tumor types (**Figure 1**), we found significant alignment gaps in all cell membrane-associated isoforms (**Figure 2F; Supplementary Figure 1D, 2F, 3D, 4D**). These observations give rise to a number of challenges, exemplified here with HER2 *(ERBB2),* which is an important growth factor receptor across a number of cancer types (**Figure 2**). For ERBB2, there were both conserved and variable regions in the ectodomain of the membrane-bound isoforms (**Figure 2F**). Should the targeted CAR epitope be located in a variable region of the ectodomain it may lead to lower treatment efficacy, or in severe cases, therapy-induced target loss. Likewise, the expression of secreted epitope harboring isoforms may cause off-tumor effects if the secreted isoforms bind to other cells, or decreased efficacy by changing the pharmacokinetics between the CAR and the cellular-bound isoform(s)^12^.

**Figure 2.**
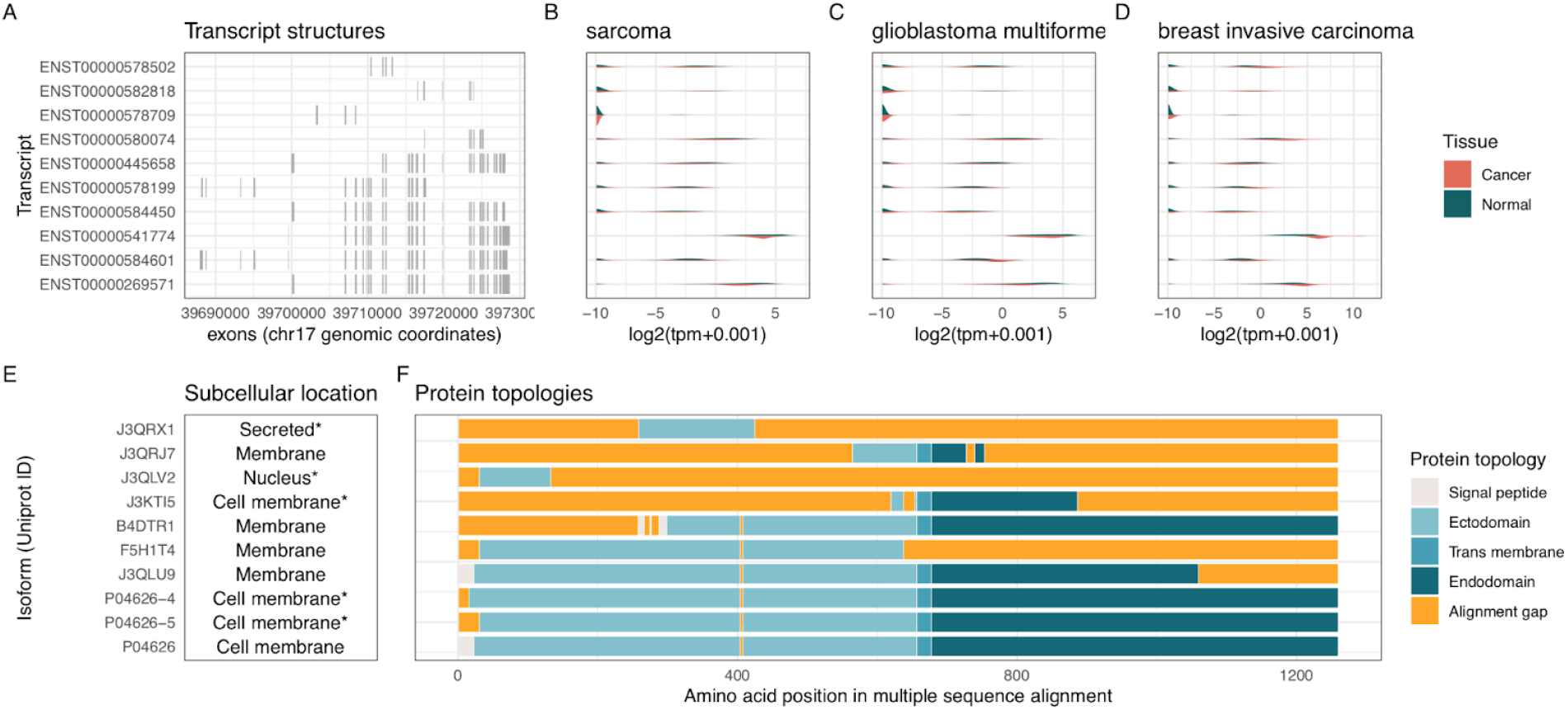
Isoform characteristics of ERBB2. (a) Genomic coordinates of exons making up the six transcripts. (b) Density distributions of transcript expression in sarcoma and all healthy tissues. (c) Density distributions of transcript expression in glioblastoma multiforme and all healthy tissues. (d) Density distributions of transcript expression in breast invasive carcinoma and all healthy tissues. (e) Subcellular location of each of the protein isoforms (predicted locations highlighted with asterisk) (f) Multiple sequence alignment of the protein isoforms with topological annotation.

We also wished to explore how sensitive newly proposed CAR targets would be in terms of variable epitope expression and secretion status should an isoform switch occur. To this end, we analyzed the targets proposed in a recently published study by MacKay *et al,* in which the authors systematically examined the expression patterns of 13,206 genes across 20 different cancers and 44 normal tissues in an effort to identify novel targets^13^. This study represents the most comprehensive of its kind but does not include analyses of the expression of different protein isoforms. In total, the authors highlight 65 potential new targets for CAR therapy. We found that 51 of these target genes express more than one protein isoform in at least one of the 20 cancers, and none of these target isoforms have completely identical ectodomains. Additionally, out of the 65 targets, 34 expresses isoforms with different subcellular locations, and 27 expresses isoforms that are potentially secreted (**Supplementary Table 1**).

Solid tumors have largely been refractory to CAR T cell therapy^14^. Here we show that the targetability of most of the currently used and proposed antigens for CAR therapy against solid tumor targets may be negatively affected by isoform switching, as has been observed with current CD19-specific CAR therapies^6–8^. Our analyses highlight the importance of considering target expression beyond the expression of the canonical protein isoform, as isoforms can have very different characteristics in terms of expression, targetable epitopes, cellular location, and secretion status.

In order to overcome this challenge, bispecific CARs could be utilized, as has been explored using an HER2/MUC1 bispecific CAR^15^, although it will still be important to consider alle expressed isoforms of these targets. Indeed both of these targets have a high number of isoforms, as shown in our analysis above. Another approach could be to promote specific isoform expression through the use of HDAC inhibitors or DNA-demethylating therapy. Here we propose another possible solution, namely by specifically targeting the small number of isoforms which are not lost, but enriched in cell-membrane proteins in cancer cells. These might provide unique opportunities for discovering truly cancer-specific targetable epitopes. For this, transcript-level, or even exon-level expression analyses, coupled with analyses of subcellular location and membrane topology of the individual protein isoforms they encode would be more informative for identifying cancer-specific structural epitopes, than the currently applied gene-level expression analyses. Such analyses would provide a space of potential targets, and technologies, such as targeted single-cell mass spectrometry^16^ are maturing to enable high-throughput, single-cell validation. As such, further studies into subclonal isoform expression are warranted, and may lead to the discovery of the first truly tumor-specific CAR targets.

