## Supplementary methods for "Target isoforms are an overlooked challenge and opportunity in chimeric antigen receptor cell therapy"

**Isoform switch analysis**

Isoform switches from TCGA along with their annotations were obtained from [1] as described in [2]. The number of genes was calculated as the number of distinct gene_name which had at least one isoform switch where the resulting protein sequences were distinct. Considering how small changes are enough to give rise to CAR cell targets we define protein isoforms as being distinct if the global pairwise alignment (Needleman-Wunsch alignment) differs with more than 5 amino acids. The pairwise alignment was done using the pairwiseAlignment function from the Biostrings R package using the type = 'global' argument. The number of amino acids differing between the two sequences was calculated as the number of amino acids (from both sequences) not part of the alignment.

We obtain a list of human transmembrane genes from UniProt [3] by filtering for the keyword “cell membrane” and for homo sapiens. Data was downloaded 5th of April 2019. To calculate the enrichment of isoform switches in transmembrane proteins we used R’s fisher.test function supplying the overlaps in gene_names.

We analyzed the identified isoform switches for differences in signal peptides (as predicted by SignalP 4.0 [4]), membrane assications (as predicted by DeepLoc [5]) and ectodomain amino acids (as predicted by TopCons2 [6]). Ectodomain amino acids are defined as “o” from “TOPCONS predicted topology” from isoforms where at least one other amino acid was predicted as not “o” (to avoid analysing secreted isoforms). An amino acid was defined as lost/gain if it was not part of the pairwise alignment described above.

**Analysis of transcript structure, tissue expression, isoform ectodomains, and subcellular locations**

Transcript level expression data was downloaded for TCGA cancer samples and GTEx healthy tissue samples from the Xena Functional Genomics Explorer [7] (www.xenabrowser.net, dataset ID “TcgaTargetGtex_rsem_isoform_tpm”). Genomic coordinates for all transcripts were retrieved from GENCODE (v36) [8], and analyzed using the GenomicRanges package for R [9].

All protein information with experimental evidence was retrieved from UniProt with the identifiers listed in **Figure 2** and **Figure 3**. Where experimental evidence of protein features was lacking, subcellular location was predicted using DeepLoc [5], signal peptides were predicted using SignalP [4], and protein topology of cell membrane spanning proteins was predicted using Topcons2 [6]. Multiple sequence alignment of protein isoform sequences was done using Clustal Omega [10]

**References**

1. Vitting-Seerup K, Sandelin A. The landscape of isoform switches in human cancers. Mol

Cancer Res. 2017;15:1206–20.

### Supplementary Figures

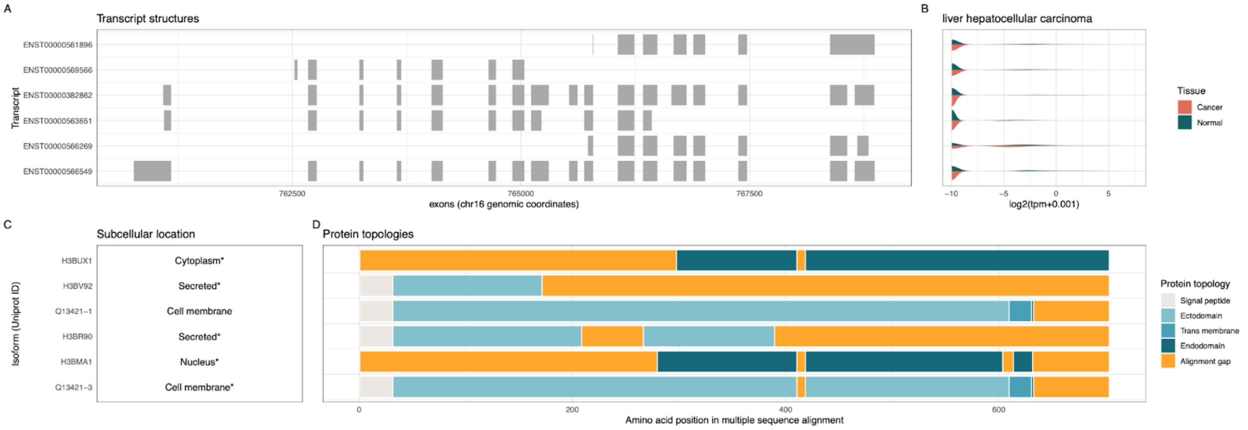


**Supplementary Figure 1**. Isoform characteristics of MSLN. (a) Genomic coordinates of exons making up the six transcripts. (b) Density distributions of transcript expression in pancreatic adenocarcinoma and all healthy tissues. (c) Subcellular location of each of the protein isoforms (e) Multiple sequence alignment of the protein isoforms with topological annotation.

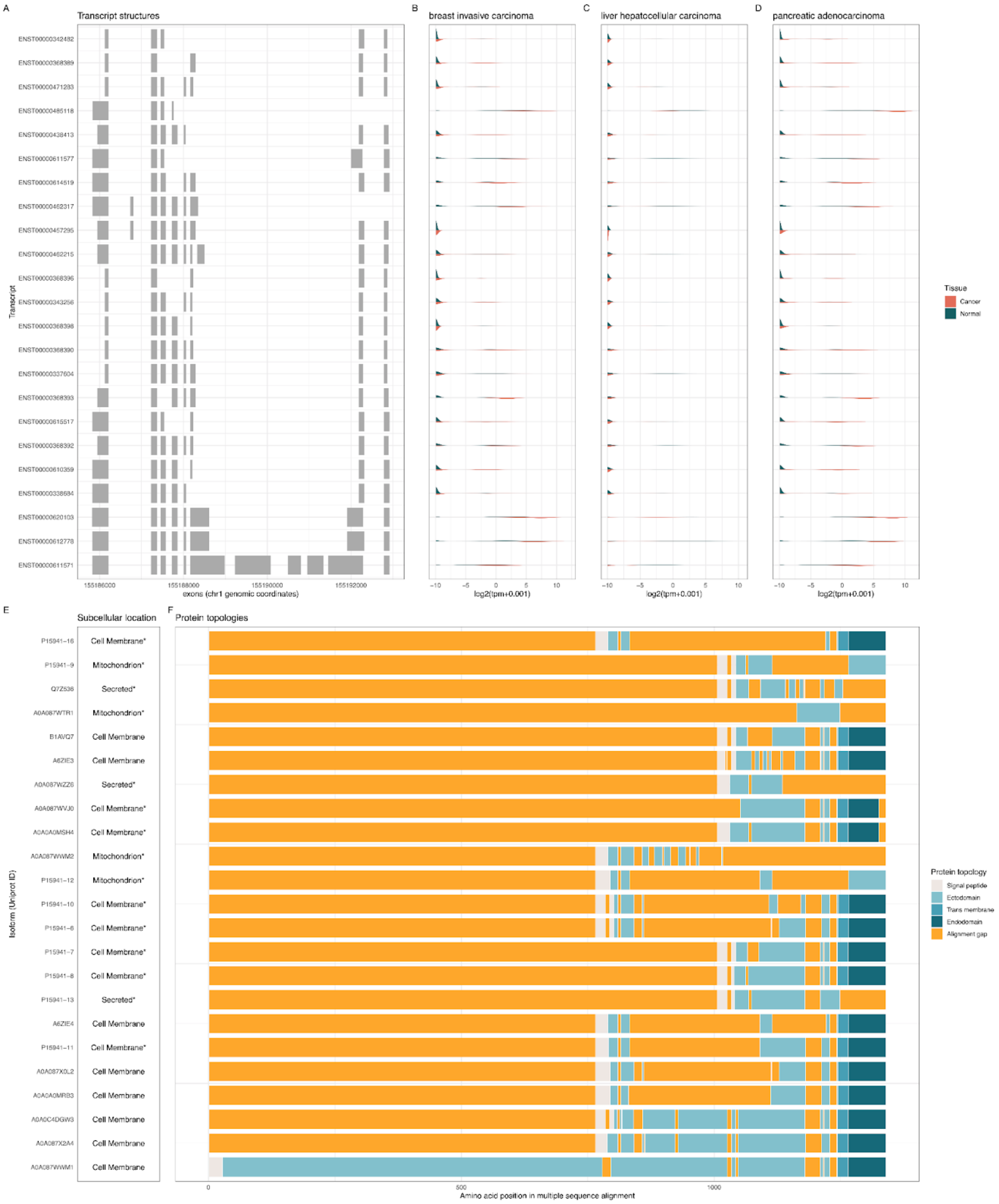


**Supplementary Figure 2**. Isoform characteristics of MUC1. (a) Genomic coordinates of exons making up the six transcripts. (b) Density distributions of transcript expression in breast invasive carcinoma and all healthy tissues. (c) Density distributions of transcript expression in liver hepatocellular carcinoma and all healthy tissues. (d) Density distributions of transcript expression in pancreatic adenocarcinoma and all healthy tissues. (e) Subcellular location of each of the protein isoforms (f) Multiple sequence alignment of the protein isoforms with topological annotation.

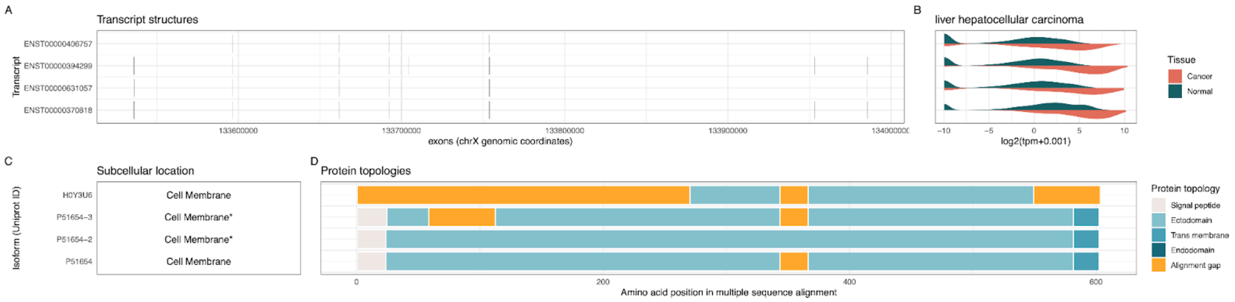

**Supplementary Figure 3**. Isoform characteristics of GPC3. (a) Genomic coordinates of exons making up the six transcripts. (b) Density distributions of transcript expression in liver hepatocellular carcinoma and all healthy tissues. (c) Subcellular location of each of the protein isoforms (e) Multiple sequence alignment of the protein isoforms with topological annotation.

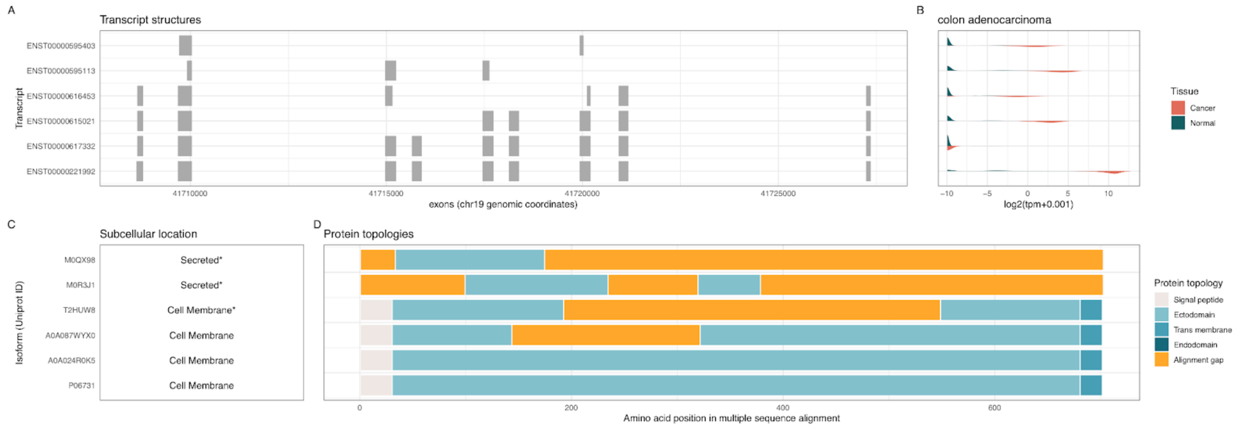


**Supplementary Figure 4**. Isoform characteristics of CEACAM5. (a) Genomic coordinates of exons making up the six transcripts. (b) Density distributions of transcript expression in colon adenocarcinoma and all healthy tissues. (c) Subcellular location of each of the protein isoforms (e) Multiple sequence alignment of the protein isoforms with topological annotation.

### Supplementary Tables

**Supplementary Table 1**: Molecular features relevant for CAR targets for each of the 65 targets suggested for CAR therapy by MacKay et al. Predicted values are denoted with and asterisk (*)

| Gene symbol | ENST | UniProt ID | Subcellular location | Ectodomain size (AA) | Signal peptide |
|---|---|---|---|---|---|
| ADGRV1 | ENST00000405460 | Q8WXG9-1 | Cell membrane | 5923 | Yes |
| ADGRV1 | ENST00000640815 | A0A1W2PQK7 | Cell membrane* | 38* | no* |
| ADGRV1 | ENST00000425867 | A0A1X7SBU6 | Cell membrane* | 2277* | no* |
| ADGRV1 | ENST00000640403 | A0A1W2PRC7 | Cytoplasm* | NA | no* |
| ADGRV1 | ENST00000638975 | A0A1W2PRR5 | Cytoplasm* | NA | no* |
| ADGRV1 | ENST00000639821 | A0A1W2PQP9 | Endoplasmic reticulum* | NA | no* |
| ADGRV1 | ENST00000639707 ENST00000640369 | A0A1W2PQU5 | Endoplasmic reticulum* | NA | no* |
| ADGRV1 | ENST00000507314 | A0A1W2PP32 | Extracellular* | NA | no* |
| ADGRV1 | ENST00000508842 | D6RIF0 | Lysosome/Vacuole* | NA | Yes |
| ALPG | ENST00000295453 | P10696 | Cell membrane | 488* | Yes |
| ALPP | ENST00000392027 | P05187 | Cell membrane | 491* | Yes |

| | | | | | |
|---|---|---|---|---|---|
| ATP6V0A4 | ENST00000310018 ENST00000353492 ENST00000393054 ENST00000645515 | Q9HBG4 | Membrane | NA | no* |
| ATP6V0A4 | ENST00000644341 | A0A2R8Y6X9 | Membrane | NA | no* |
| ATP6V0A4 | ENST00000478480 | A0A2R8YE46 | Membrane | NA | no* |
| ATP6V0A4 | ENST00000483139 | A0A2R8YGN5 | Membrane | NA | no* |
| BPI | ENST00000642449 | A0A2R8YDF1 | Extracellular | 456* | Yes |
| BPI | ENST00000262865 | P17213 | Extracellular | 457* | Yes |
| BPI | ENST00000417318 | H0Y738 | Extracellular | 286* | yes* |
| C3AR1 | ENST00000307637 | Q16581 | Cell membrane | 1520 | no* |
| C3AR1 | ENST00000546241 | F5GZE6 | Cell membrane | 41* | no* |
| CACNG3 | ENST00000005284 | O60359 | Membrane | NA | no* |
| CCNB3 | ENST00000276014 | Q8WWL7 | Nucleus | NA | no* |
| CCNB3 | ENST00000376042 | Q8WWL7 | Nucleus | NA | no* |
| CCNB3 | ENST00000376038 | Q8WWL7-2 | Nucleus* | NA | no* |

| | | | | | |
|---|---|---|---|---|---|
| CCNB3 | ENST00000348603 | Q8WWL7-2 | Nucleus* | NA | no* |
| CCR2 | ENST00000292301 | P41597 | Cell membrane | 98 | no* |
| CCR2 | ENST00000421659 | E9PH76 | Cell membrane | 67* | no* |
| CCR2 | ENST00000400888 | P41597 | Cell membrane | 209* | no* |
| CCR2 | ENST00000445132 | P41597-2 | Cell membrane* | 194* | no* |
| CCR8 | ENST00000326306 | P51685 | Cell membrane | 94 | no* |
| CCR8 | ENST00000414803 | C9JIP9 | Cytoplasm* | NA | no* |
| CD2 | ENST00000369478 | P06729 | Cell membrane | 284 | Yes |
| CD2 | ENST00000369477 | Q5JVN7 | Cell membrane | 112* | Yes |
| CD27 | ENST00000266557 | P26842 | Cell membrane | 171 | Yes |
| CD37 | ENST00000323906 | P11049 | Cell membrane | 149 | no* |
| CD37 | ENST00000426897 | P11049-2 | Cell membrane* | 131* | no* |
| CD37 | ENST00000598095 | P11049-3 | Endoplasmic reticulum* | NA | no* |
| CD37 | ENST00000597602 | M0R083 | Extracellular* | NA | no* |
| CD37 | ENST00000535669 | B7ZAN3 | Membrane | NA | no* |
| CD37 | ENST00000595660 | M0QZ33 | Membrane | NA | no* |

| | | | | | |
|---|---|---|---|---|---|
| CD37 | ENST00000594743 | M0QZP6 | Membrane | NA | no* |
| CD3D | ENST00000300692 | P04234-1 | Cell membrane | 83 | Yes |
| CD3D | ENST00000392884 | P04234-2 | Extracellular* | NA | yes* |
| CD3D | ENST00000529594 | E9PMT5 | Membrane | NA | Yes |
| CD3D | ENST00000534687 | H0YE18 | Membrane | NA | no* |
| CD3G | ENST00000532917 | P09693 | Cell membrane | 93 | Yes |
| CD3G | ENST00000392883 | A8MUH3 | Membrane | NA | no* |
| CD3G | ENST00000528540 | A0A3B3IUD8 | Mitochondrion* | NA | no* |
| CD72 | ENST00000259633 | P21854 | Cell membrane | 242 | no* |
| CD72 | ENST00000612238 | A0A087X0Y7 | Cell membrane* | 244* | no* |
| CD72 | ENST00000378431 | Q5T4Q8 | Cell membrane* | 4* | no* |
| CD72 | ENST00000396757 | A0A6E1WA84 | Endoplasmic reticulum* | NA | no* |
| CD72 | ENST00000378430 | Q5T4Q7 | Plastid* | NA | no* |
| CD79A | ENST00000221972 | P11912-1 | Cell membrane | 110 | Yes |
| CD79A | ENST00000597454 | M0QX61 | Cell membrane | 228* | yes* |
| CD79A | ENST00000444740 | P11912-2 | Cell membrane* | 76* | yes* |

| | | | | | |
|---|---|---|---|---|---|
| CD79B | ENST00000006750 | P40259-1 | Cell membrane | 130 | Yes |
| CD79B | ENST00000392795 | P40259-3 | Cell membrane* | 131* | yes* |
| CD79B | ENST00000349817 | P40259-2 | Endoplasmic reticulum* | NA | yes* |
| CD83 | ENST00000379153 | Q01151 | Cell membrane | 124 | Yes |
| CD83 | ENST00000612003 | A0A087WX61 | Endoplasmic reticulum* | NA | no* |
| CEACAM3 | ENST00000344550 | P40198-3 | Cell membrane | 120 | Yes |
| CEACAM3 | ENST00000357396 | P40198 | Cell membrane* | 152* | yes* |
| CEACAM3 | ENST00000630848 | P40198-3 | Cell membrane* | 121* | yes* |
| CEACAM3 | ENST00000596544 | M0QXR5 | Cytoplasm* | NA | no* |
| CEACAM3 | ENST00000415495 | P40198-2 | Extracellular* | NA | yes* |
| CEACAM8 | ENST00000244336 | P31997 | Cell membrane | 297* | Yes |
| CEACAM8 | ENST00000599005 | M0R1X3 | Extracellular* | NA | no* |
| CELSR3 | ENST00000164024 | Q9NYQ7-1 | Cell membrane | 2524 | Yes |
| DCLK2 | ENST00000296550 | Q8N568 | Cytoplasm | NA | no* |
| DCLK2 | ENST00000635524 | A0A0U1RR70 | Cytoplasm* | NA | no* |
| DCLK2 | ENST00000506325 | Q8N568-2 | Cytoplasm* | NA | no* |

| | | | | | |
|---|---|---|---|---|---|
| DCLK2 | ENST00000302176 | Q8N568-3 | Cytoplasm* | NA | no* |
| DYRK4 | ENST00000010132 | Q9NR20 | Cytoplasm | NA | no* |
| DYRK4 | ENST00000540757 | Q9NR20 | Cytoplasm* | NA | no* |
| DYRK4 | ENST00000543431 | A0A0A0MTH5 | Nucleus* | NA | no* |
| DYRK4 | ENST00000544671 | H0YFI4 | Nucleus* | NA | no* |
| FASLG | ENST00000340030 | P48023-2 | Cell membrane | 178 | no* |
| FASLG | ENST00000367721 | P48023 | Cell membrane* | 180* | no* |
| FCRL1 | ENST00000358292 | Q96LA6-3 | Cell membrane | 290 | Yes |
| FCRL1 | ENST00000368176 | Q96LA6 | Cell membrane* | 287* | yes* |
| FCRL1 | ENST00000491942 | Q96LA6-2 | Cell membrane* | 287* | yes* |
| FCRLA | ENST00000236938 | Q7L513 | Cytoplasm | NA | Yes |
| FCRLA | ENST00000367959 | Q7L513-10 | Endoplasmic reticulum* | NA | no* |
| FCRLA | ENST00000546024 | Q7L513-12 | Endoplasmic reticulum* | NA | no* |
| FCRLA | ENST00000674323 | Q7L513-9 | Endoplasmic reticulum* | NA | no* |
| FCRLA | ENST00000367950 | Q5VXA5 | Extracellular* | NA | Yes |
| FCRLA | ENST00000367949 | Q7L513-13 | Extracellular* | NA | no* |

| | | | | | |
|---|---|---|---|---|---|
| FCRLA | ENST00000350710 | Q7L513-15 | Extracellular* | NA | no* |
| FCRLA | ENST00000367953 | Q7L513-2 | Extracellular* | NA | yes* |
| FCRLA | ENST00000309691 | Q7L513-3 | Extracellular* | NA | yes* |
| FCRLA | ENST00000294796 | Q7L513-4 | Extracellular* | NA | no* |
| FCRLA | ENST00000367957 | Q7L513-5 | Extracellular* | NA | no* |
| FCRLA | ENST00000349527 | Q7L513-8 | Extracellular* | NA | yes* |
| FCRLA | ENST00000540521 | Q7L513-11 | Golgi apparatus* | NA | no* |
| FCRLA | ENST00000674251 | Q7L513-14 | Golgi apparatus* | NA | no* |
| GLRB | ENST00000264428 | P48167 | Cell membrane | 255 | Yes |
| GLRB | ENST00000509282 | P48167 | Cell membrane | 308* | yes* |
| GLRB | ENST00000541722 | P48167-2 | Cell membrane* | 245* | yes* |
| GLRB | ENST00000512619 | D6RD86 | Extracellular* | NA | Yes |
| GRM2 | ENST00000395052 | Q14416 | Cell membrane | 591 | Yes |
| GRM2 | ENST00000442933 | C9JL63 | Cell membrane | 496* | Yes |
| GRM2 | ENST00000419928 | C9JD41 | Extracellular* | NA | Yes |
| GYPA | ENST00000324022 | P02724-3 | Cell membrane | 71 | Yes |

| | | | | | |
|---|---|---|---|---|---|
| GYPA | ENST00000360771 | A0A0C4DFT7 | Cell membrane* | 69* | Yes |
| GYPA | ENST00000504786 | E7EQF3 | Cell membrane* | 38* | Yes |
| GYPA | ENST00000512064 | E9PD10 | Cell membrane* | 56* | Yes |
| GYPA | ENST00000503627 | E9PH25 | Cell membrane* | 25* | Yes |
| GYPA | ENST00000641688 | A0A0C4DFT7 | Cell membrane* | 69* | yes* |
| GYPA | ENST00000643148 | K9JIK7 | Cell membrane* | 12* | no* |
| GYPA | ENST00000512789 | Q13030 | Cell membrane* | 25* | no* |
| GYPA | ENST00000642295 | A0A2R8Y7F9 | Endoplasmic reticulum* | NA | Yes |
| GYPA | ENST00000616983 | A0A087WU29 | Endoplasmic reticulum* | NA | no* |
| GYPA | ENST00000535709 | A0A087WU29 | Endoplasmic reticulum* | NA | no* |
| GYPA | ENST00000642738 | K9JI14 | Endoplasmic reticulum* | NA | yes* |
| GYPA | ENST00000642713 | P02724-2 | Endoplasmic reticulum* | NA | no* |
| HEPACAM | ENST00000298251 | Q14CZ8-1 | Cytoplasm | NA | Yes |
| HERC5 | ENST00000264350 | Q9UII4 | Cytoplasm | NA | no* |
| HERC5 | ENST00000508159 | E9PBL0 | Cytoplasm* | NA | no* |
| HIST1H1T | ENST00000338379 | P22492 | Nucleus | NA | no* |

| | | | | | |
|---|---|---|---|---|---|
| HMMR | ENST0000035 3866 | O75330-2 | Cell membrane | 709* | no* |
| HMMR | ENST0000035 8715 | O75330 | Golgi apparatus* | NA | no* |
| HMMR | ENST0000039 3915 | O75330-3 | Golgi apparatus* | NA | no* |
| HMMR | ENST0000043 2118 | O75330-4 | Golgi apparatus* | NA | no* |
| HMMR | ENST0000052 0345 | E5RI30 | Nucleus* | NA | no* |
| HMMR | ENST0000052 2094 | E5RIH2 | Nucleus* | NA | no* |
| IZUMO4 | ENST0000058 8003 | A0A087X2 21 | Cytoplasm* | NA | no* |
| IZUMO4 | ENST0000062 0263 | A0A087X1 M1 | Extracellular* | NA | Yes |
| IZUMO4 | ENST0000039 5296 | A0A0A0M S61 | Extracellular* | NA | Yes |
| IZUMO4 | ENST0000039 5307 | Q1ZYL8-2 | Extracellular* | NA | yes* |
| IZUMO4 | ENST0000061 0800 | A0A087X1 60 | Nucleus* | NA | no* |
| IZUMO4 | ENST0000039 5301 | Q1ZYL8 | Extracellular | 211* | Yes |
| KCNN4 | ENST0000064 8319 | O15554 | Cell membrane | 104* | no* |
| KCNN4 | ENST0000061 5047 | D1MQ08 | Membrane | NA | no* |
| KCNN4 | ENST0000059 8836 | M0R1J0 | Membrane | NA | no* |
| KCNN4 | ENST0000060 0909 | M0QZ70 | Nucleus* | NA | no* |

| | | | | | |
|---|---|---|---|---|---|
| KLK2 | ENST0000039 1810 | P20151-4 | Cytoplasm* | NA | no* |
| KLK2 | ENST0000060 0690 | M0QXQ7 | Extracellular* | NA | Yes |
| KLK2 | ENST0000032 5321 | P20151 | Extracellular* | NA | Yes |
| KLK2 | ENST0000035 8049 | P20151-2 | Extracellular* | NA | yes* |
| KLK2 | ENST0000059 7439 | P20151-3 | Extracellular* | NA | yes* |
| KLK2 | ENST0000059 9568 | M0R0M4 | Nucleus* | NA | no* |
| KLK2 | ENST0000059 3493 | M0R2W5 | Nucleus* | NA | no* |
| KLK3 | ENST0000059 7286 | M0QZF9 | Extracellular* | NA | Yes |
| KLK3 | ENST0000059 8145 | M0R1F0 | Extracellular* | NA | Yes |
| KLK3 | ENST0000060 1503 | M0R1Z7 | Extracellular* | NA | Yes |
| KLK3 | ENST0000059 7483 | M0R294 | Extracellular* | NA | Yes |
| KLK3 | ENST0000061 7027 | Q8NCW4 | Extracellular* | NA | Yes |
| KLK3 | ENST0000036 0617 | P07288-2 | Extracellular* | NA | yes* |
| KLK3 | ENST0000059 5952 | P07288-3 | Extracellular* | NA | yes* |
| KLK3 | ENST0000059 3997 | P07288-5 | Extracellular* | NA | yes* |
| KLK3 | ENST0000032 6003 | P07288 | Extracellular | 239* | Yes |

| | | | | | |
|---|---|---|---|---|---|
| LILRA4 | ENST00000291759 | P59901 | Cell membrane | 422 | Yes |
| MS4A8 | ENST00000529752 | E9PQE1 | Cell membrane* | 24* | no* |
| MS4A8 | ENST00000525458 | H0YD94 | Cell membrane* | 29* | no* |
| MS4A8 | ENST00000300226 | Q9BY19 | Membrane | NA | no* |
| MUC12 | ENST00000536621 | Q9UKN1-2 | Cell membrane* | 5213* | yes* |
| MUC12 | ENST00000379442 | Q9UKN1-1 | Membrane | NA | Yes |
| NANOG | ENST00000541267 | F5GZI2 | Nucleus | NA | no* |
| NANOG | ENST00000229307 | Q9H9S0 | Nucleus | NA | no* |
| NANOG | ENST00000526286 | Q9H9S0-2 | Nucleus* | NA | no* |
| NBPF3 | ENST00000318249 | Q9H094 | Cytoplasm | NA | no* |
| NBPF3 | ENST00000619554 | Q9H094-2 | Cytoplasm* | NA | no* |
| NBPF3 | ENST00000342104 | Q9H094-3 | Cytoplasm* | NA | no* |
| NBPF3 | ENST00000454000 | Q9H094-5 | Cytoplasm* | NA | no* |
| NGB | ENST00000298352; | Q9NPG2 | Cytoplasm | NA | no* |
| OTUB2 | ENST00000203664 | Q96DC9 | Cytoplasm* | NA | no* |
| OTUB2 | ENST00000553723 | Q96DC9-2 | Mitochondrion* | NA | no* |

| | | | | | |
|---|---|---|---|---|---|
| PSG1 | ENST00000595930 | M0QY44 | Cytoplasm* | NA | no* |
| PSG1 | ENST00000597058 | M0QZQ1 | Cytoplasm* | NA | no* |
| PSG1 | ENST00000595124 | M0R235 | Endoplasmic reticulum* | NA | Yes |
| PSG1 | ENST00000595356 | P11464-3 | Endoplasmic reticulum* | NA | yes* |
| PSG1 | ENST00000403380 | G5E9F7 | Extracellular* | NA | Yes |
| PSG1 | ENST00000436291 | P11464 | Extracellular* | NA | yes* |
| PSG1 | ENST00000312439 | P11464-2 | Extracellular* | NA | yes* |
| PSG1 | ENST00000244296 | P11464-4 | Extracellular | 396* | Yes |
| RBMXL2 | ENST00000306904 | O75526 | Nucleus | NA | no* |
| RBMXL3 | ENST00000424776 | Q8N7X1 | Nucleus* | NA | no* |
| SIGLEC8 | ENST00000430817 | C9JT30 | Cell membrane* | 251* | Yes |
| SIGLEC8 | ENST00000340550 | Q9NYZ4-2 | Cell membrane* | 267* | yes* |
| SIGLEC8 | ENST00000321424 | Q9NYZ4-1 | Membrane | NA | Yes |
| SLC13A5 | ENST00000293800 | Q86YT5-3 | Cell membrane | 98* | no* |
| SLC13A5 | ENST00000433363 | Q86YT5 | Cell membrane* | 105* | no* |
| SLC13A5 | ENST00000573648 | Q86YT5-2 | Cell membrane* | 103* | no* |

| | | | | | |
|---|---|---|---|---|---|
| SLC13A5 | ENST00000381074 | Q86YT5-4 | Cell membrane* | 103* | no* |
| SLC13A5 | ENST00000570687 | I3L2Y7 | Membrane | NA | Yes |
| SLC13A5 | ENST00000572352 | I3L4X6 | Membrane | NA | no* |
| SLC22A12 | ENST00000336464 | Q96S37-4 | Cell membrane | 51* | no* |
| SLC22A12 | ENST00000377574 | Q96S37 | Cell membrane* | 141* | no* |
| SLC22A12 | ENST00000377567 | Q96S37-2 | Cell membrane* | 134* | no* |
| SLC22A12 | ENST00000377572 | Q96S37-2 | Cell membrane* | 134* | no* |
| SLC22A12 | ENST00000473690 | Q96S37-3 | Endoplasmic reticulum* | NA | no* |
| SLC2A14 | ENST00000340749 | Q8TDB8-2 | Cell membrane | 85 | no* |
| SLC2A14 | ENST00000431042 | Q8TDB8-2 | Cell membrane | 69* | no* |
| SLC2A14 | ENST00000616981 | Q8TDB8 | Cell membrane* | 69* | no* |
| SLC2A14 | ENST00000543909 | Q8TDB8 | Cell membrane* | 69* | no* |
| SLC2A14 | ENST00000396589 | Q8TDB8 | Cell membrane* | 69* | no* |
| SLC2A14 | ENST00000542505 | Q8TDB8-3 | Cell membrane* | 5* | no* |
| SLC2A14 | ENST00000542546 | Q8TDB8-4 | Cell membrane* | 34* | no* |
| SLC2A14 | ENST00000535295 | Q8TDB8-4 | Cell membrane* | 34* | no* |

| | | | | | |
|---|---|---|---|---|---|
| SLC2A14 | ENST00000539924 | Q8TDB8-5 | Cell membrane* | 68* | no* |
| SLC2A14 | ENST00000535344 | F5H5Q3 | Endoplasmic reticulum* | NA | no* |
| SLC2A14 | ENST00000535587 | F5H3J0 | Extracellular* | NA | no* |
| SLC2A14 | ENST00000539234 | F5H565 | Extracellular* | NA | no* |
| SLC2A14 | ENST00000546234 | F5GXP8 | Lysosome/Vacuole* | NA | no* |
| SLC2A14 | ENST00000542782 | F5H076 | Lysosome/Vacuole* | NA | no* |
| SLC2A14 | ENST00000535383 | F5H5V7 | Mitochondrion* | NA | no* |
| SLC2A14 | ENST00000542916 | F5H7N2 | Mitochondrion* | NA | no* |
| SLC2A14 | ENST00000537557 | F5GXP7 | Plastid* | NA | no* |
| SLC2A14 | ENST00000535266 | F5H6F6 | Plastid* | NA | no* |
| SLC34A1 | ENST00000324417 | Q06495 | Cell membrane | 213 | no* |
| SLC34A1 | ENST00000504577 | D6RCE5 | Cell membrane | 109* | no* |
| SLC34A1 | ENST00000512593 | Q06495-2 | Plastid* | NA | no* |
| SLC45A3 | ENST00000367145 | Q96JT2 | Membrane | NA | no* |
| SLC46A2 | ENST00000374228 | Q9BY10 | Cell membrane | 76 | no* |
| SLC4A1 | ENST00000262418 | P02730 | Cell membrane | 82 | no* |

| | | | | | |
|---|---|---|---|---|---|
| SLC4A1 | ENST00000399246 | A0A0A0MS98 | Membrane | NA | no* |
| SLC4A1 | ENST00000631130 | V9H0V9 | Mitochondrion* | NA | no* |
| SLC6A11 | ENST00000454147 | P48066-2 | Cell membrane* | 33* | no* |
| SLC6A11 | ENST00000254488 | P48066 | Membrane | NA | no* |
| SLC7A3 | ENST00000298085 ENST00000374299 | Q8WY07 | Cell membrane | 90 | no* |
| SPATA19 | ENST00000299140 | Q7Z5L4 | Mitocondria | NA | no* |
| SV2C | ENST00000502798 | Q496J9 | Cytoplasm | NA | no* |
| SV2C | ENST00000322285 | B3KT41 | Cytoplasm | NA | no* |
| TNFRSF13C | ENST00000291232 | Q96RJ3 | Membrane | NA | no* |
| TSPAN16 | ENST00000590327 | Q9UKR8-2 | Cell membrane* | 127* | no* |
| TSPAN16 | ENST00000592955 | Q9UKR8-3 | Cell membrane* | 105* | no* |
| TSPAN16 | ENST00000621731 | Q9UKR8-4 | Lysosome/Vacuole* | NA | no* |
| TSPAN16 | ENST00000337994 | Q9UKR8-4 | Lysosome/Vacuole* | NA | no* |
| TSPAN16 | ENST00000316737 | Q9UKR8 | Membrane | NA | no* |
| UPK1A | ENST00000617999 | O00322 | Cell membrane* | 144* | no* |

| | | | | | |
|---|---|---|---|---|---|
| UPK1A | ENST0000037 9013 | O00322-2 | Cell membrane* | 185* | no* |
| UPK1A | ENST0000061 6789 | O00322-2 | Cell membrane* | 185* | no* |
| UPK1A | ENST0000022 2275 | O00322 | Membrane | NA | no* |
| UPK3B | ENST0000025 7632 | Q9BT76-1 | Cell membrane | 0 | Yes |
| UPK3B | ENST0000033 4348 | Q9BT76-3 | Cell membrane* | 170* | yes* |
| UPK3B | ENST0000039 4849 | Q9BT76-2 | Extracellular* | NA | yes* |
| ZP2 | ENST0000057 4091 | Q05996-2 | Cell membrane* | 664* | no* |
| ZP2 | ENST0000064 0487 | Q05996-2 | Cell membrane* | 664* | no* |
| ZP2 | ENST0000057 4002 | Q05996 | Extracellular | 677 | Yes |
| ZP2 | ENST0000063 8300 | Q05996 | Extracellular | 673* | yes* |
